# Threshold responses of multi-trophic freshwater communities to browning and eutrophication

**DOI:** 10.64898/2026.04.22.719940

**Authors:** Maude Lachapelle, Irene Gregory-Eaves, Susanne Kraemer, Marc Amyot, Marie-Ève Monchamp, Marie-Pier Hébert, Michelle Gros, Zofia E. Taranu

**Affiliations:** Aquatic Contaminant Research Division, Environment and Climate Change Canada; Groupe de Recherche Interuniversitaire en Limnologie (GRIL); Biology Department, McGill University; Département des Sciences Biologiques, Université de Montréal; Département des sciences de l’environnement, Université du Québec à Trois-Rivières

**Keywords:** Browning, Eutrophication, Biodiversity, Multi-trophic (bacterioplankton, phytoplankton, zooplankton), eDNA, Thresholds

## Abstract

Browning and eutrophication strongly influence aquatic ecosystems by altering nutrient dynamics, light availability, and food web structure. To investigate their combined effects on aquatic communities, we conducted a nine-week mesocosm experiment in a clear-water north-temperate lake, crossing dissolved organic carbon (DOC) and total phosphorus plus total nitrogen (TP+TN) enrichment treatments. Multi-trophic plankton communities (bacterioplankton, phytoplankton, and zooplankton) were monitored over time using environmental DNA (eDNA) marker gene amplicon sequencing. Beta-diversity analyses highlighted temporal and treatment-driven community restructuring, while PERMANOVA and Principal Response Curve analyses identified the treatments and taxa driving these changes. Our results show that elevated DOC favoured taxa associated with the microbial loop, while nutrient enrichment and lower DOC promoted the green pathway. Threshold responses across trophic levels were observed at 5–7 mg L⁻¹ DOC and 30–70 μg L⁻¹ TP, marking the levels at which compositional shifts propagated through the food web. Overall, this study demonstrated how aquatic communities respond dynamically to browning and nutrient enrichment, offering insight into the mechanisms shaping multi-trophic interactions under a multiple stressor scenario.

**Highlights:**

- Browning and nutrient pulses drove coordinated shifts across bacterioplankton, phytoplankton, and zooplankton.
- Temporal community succession and treatment effects were captured through beta-diversity and multivariate ordination analyses.
- Threshold responses propagated through the food web at 5–7 mg L⁻¹ DOC and 30–70 µg L⁻¹ TP.

**Scientific Significance Statement Topic:** This study provides insights into the interactive effects of browning and eutrophication on community composition shifts in freshwater ecosystems. Using a mesocosm experiment, we identified thresholds for dissolved organic carbon and total phosphorus + nitrogen concentrations that drove compositional changes across bacterial, phytoplankton, and zooplankton communities, as determined by environmental DNA amplicon sequencing data.

**Graphical Abstract:** 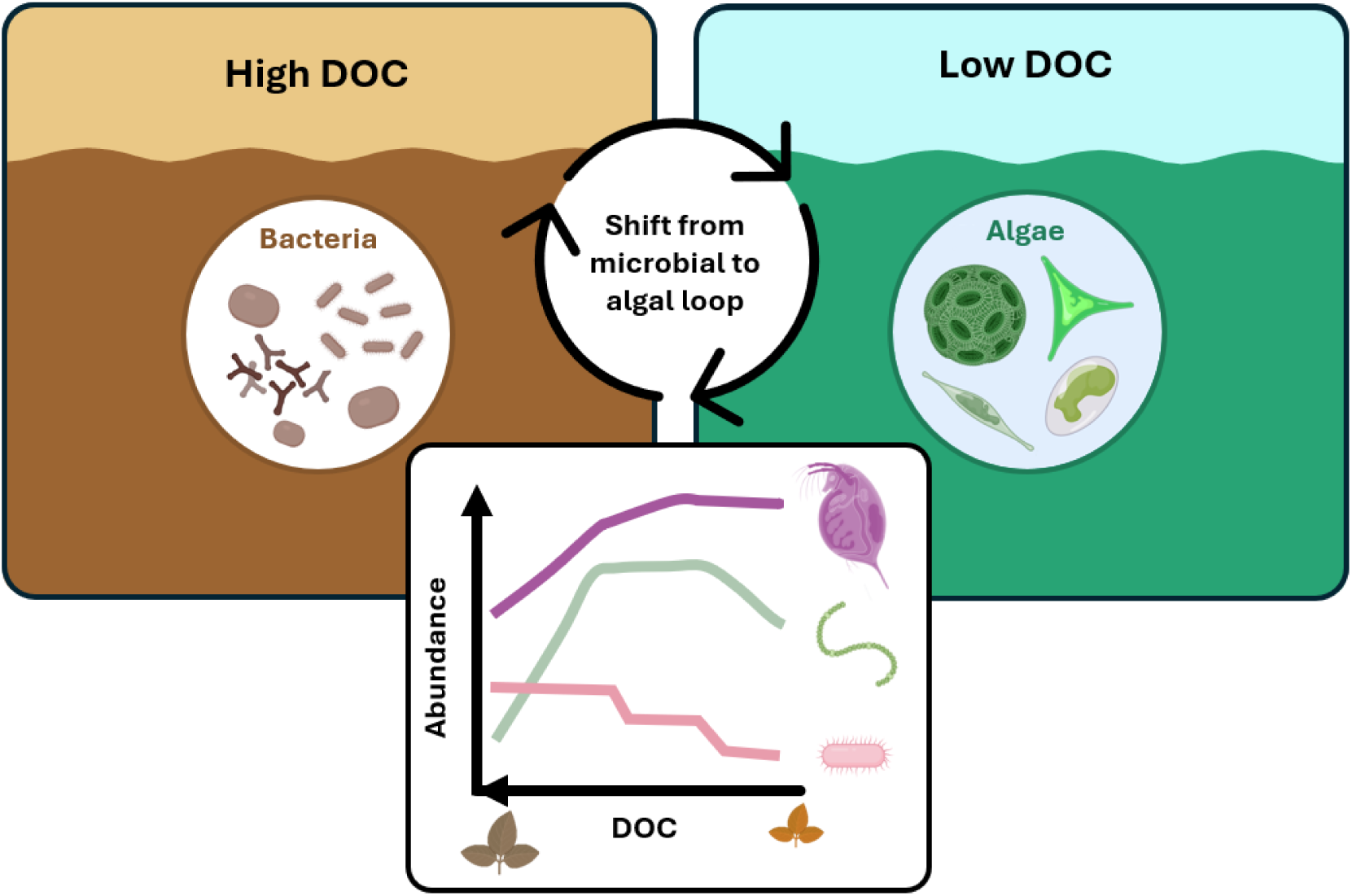

## Introduction

Globally, humans are altering biogeochemical cycles, such as carbon, nitrogen (N), and phosphorus (P), at an accelerating rate (Steffen et al., 2007; Fowler et al., 2013; Mekonnen & Hoekstra, 2018).

Anthropogenic activities have dramatically modified the carbon cycle over the past century, while the use of synthetic nitrogen fertilizers has increased nine-fold globally since the 1960s, and phosphorus fertilizers use has tripled, with further increases projected (Steffen et al., 2007; Sutton et al., 2013; Steffen et al., 2015; Waters et al., 2016). As a result, many downstream lakes are darkening, a process that can arise due to different mechanisms: some lakes are greening due to nutrient enrichment and algal blooms (Williamson, 2020; Xenopoulos et al., 2021), others are browning due to terrestrial organic matter inputs (Monteith et al., 2007; Kritzberg 2017), and some are becoming murkier (i.e., both greening and browning; Leech et al. 2018; Hayden et al. 2019). These processes modify light penetration and reshape planktonic food webs (Berggren et al. 2015), including changes in phytoplankton (e.g., increases in cyanobacteria and mixotrophic taxa; Creed et al., 2018; Farragher M.J., 2021; Senar et al., 2021; Horppila et al., 2023) and zooplankton (e.g., reliance on bacterioplankton- versus phytoplankton-based food chains; Berggren et al. 2014; Senar et al., 2019; Hébert et al., 2022; Tonin et al., 2022; Strandberg et al., 2023). Single-stressor thresholds that trigger community shifts vary considerably in the literature, making it difficult to predict how communities will respond when exposed to multiple stressors simultaneously (e.g., DOC thresholds ranging from 2-16 mg/L or from 5-9 mg/L for changes in phytoplankton and zooplankton community structures, respectively; Farragher M.J., 2021; Horppila et al., 2023; Tonin et al., 2022; Young et al., 2024). Despite previous studies suggesting dual effects of browning and eutrophication, the individual contributions of dissolved organic carbon (DOC) and nutrient (nitrogen and phosphorus) enrichment across lakes or over time remain hard to disentangle. Understanding how these drivers restructure communities across trophic levels and how changes propagate through food webs is crucial for predicting how browning and eutrophication jointly shape plankton community composition and energy pathways in freshwater ecosystems.

Our experimental study was designed to quantify the single and combined effects of browning and eutrophication on plankton community composition in a clear-water mesotrophic lake, within a background of seasonal variation. We experimentally induced browning (increased water color via DOC enrichment) and eutrophication (P and N addition) within lake mesocosms from late summer (peak water temperatures of 25 °C) to late fall (lows of 10 °C). Responses in aquatic communities were monitored through high-throughput sequencing (HTS) of environmental DNA (eDNA), targeting bacterioplankton, phytoplankton and zooplankton. We hypothesized that browning (high DOC) would favor taxa tolerant to low light conditions, including mixotrophs and heterotrophs, while eutrophication (high TP+TN) would boost primary productivity (Creed et al., 2018; Senar et al., 2019). Our work set out to: 1) test whether nutrient and DOC additions induced the anticipated shifts in communities, 2) identify potential indicator taxa of browning and eutrophication, and 3) determine the threshold concentrations that triggered changes in community composition. This design allowed us to examine how short-term responses at different trophic levels are interconnected and propagate through lake food webs.

## Materials and Methods

*Experimental setup.* We ran a nine-week mesocosm experiment from August 25^th^ to October 31^st^, 2021, in Lake Hertel, Quebec (45°32′N, 73°08′W). The site is a temperate headwater lake in eastern Canada, which stratifies over summer and is in a pristine forested catchment (Figure 1a). In total, 18 mesocosms were deployed on a floating dock off the shore of the lake (Figure 1b). Each mesocosm consisted of a 3-m deep plastic bag that was 1 m in diameter, with a capacity of approximately 2400 L of water (no sediments).

**Figure 1.**
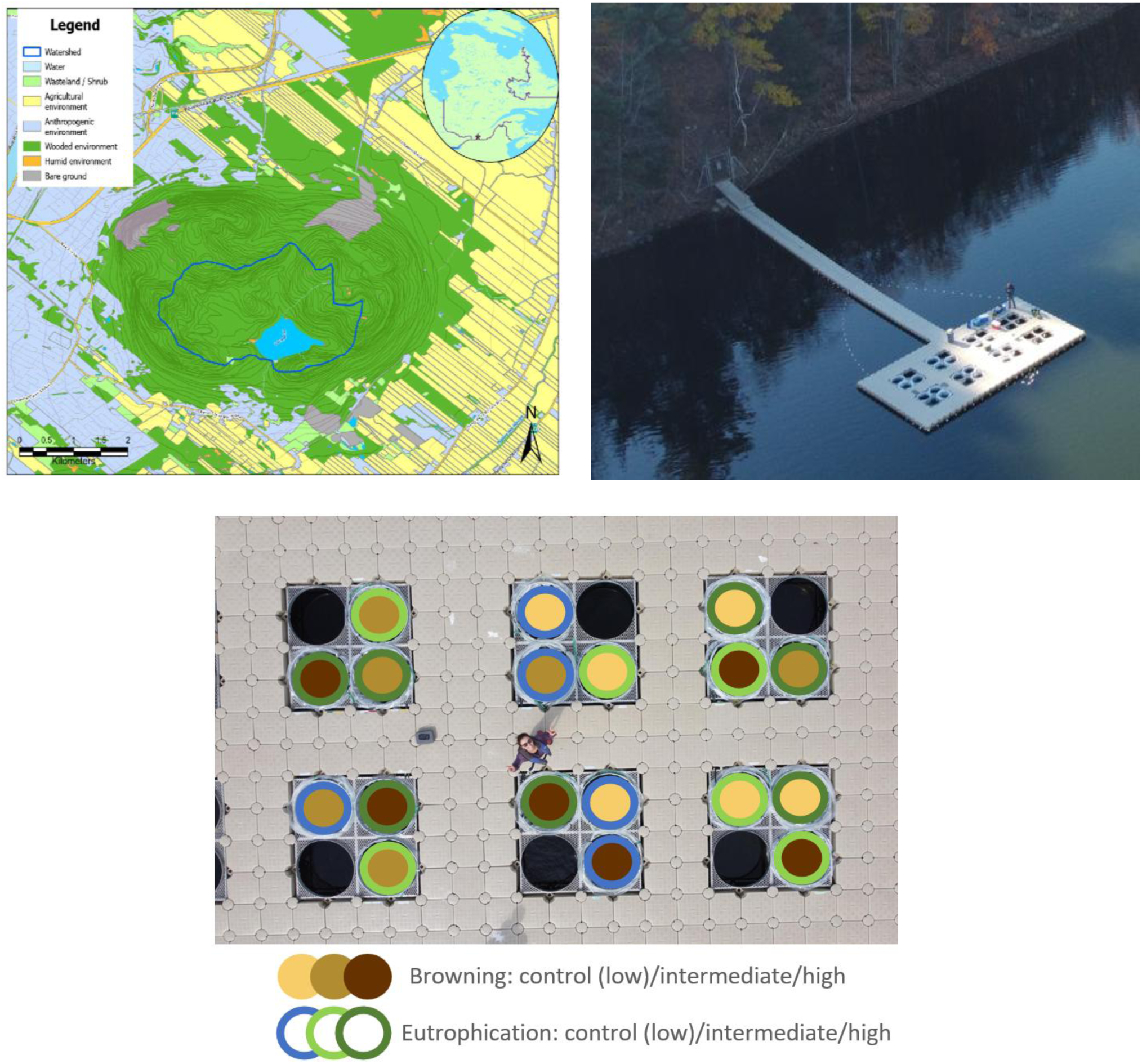
Mesocosm setup, with **a)** map of study site and its surrounding landscape, **b)** aerial photograph of the mesocosms on the floating dock setup on Lake Hertel, and **c)** diagram of the experimental treatments crossing browning (tan to brown) and eutrophication (blue to green).

Each bag was filled with lake water and organisms collected from 0.5 to 2.5 m depths.

To mimic lake eutrophication and browning, the mesocosms were treated in a duplicated semi-factorial design of low (mesotrophic lake water used as a control baseline), intermediate, or high TN+TP and DOC additions (Figure 1c). Eutrophication (TN+TP) treatments were targeted to double (intermediate) and triple (high) the baseline Lake Hertel of TN (∼0.30 mg/L) and TP (∼20 μg/L) concentrations. For the browning treatments (coloured dissolved organic matter (DOM) additions, measured as DOC), we conducted a two-week leaching incubation on the lake shore to extract and concentrate terrestrially-derived DOM from local leaf litter and topsoil (final DOC concentration in leachate inoculum ∼125 mg/L). We filtered leachates (DOM concentrates) through a 120 μm mesh to reduce the addition of particulate matter to mesocosms, and then added the filtrates to each mesocosm to reach target concentrations of 7.5 mg/L DOC for the intermediate and 10.5 mg/L DOC for the high browning treatment. The baseline DOC level of Lake Hertel (control) was at 3 mg/L.

*Sample collection and processing.* Following the treatment spikes on week 0, mesocosms were sampled at 1-week (physico-chemistry), 2-week (DOC, TP, TN), and 4-week (eDNA) intervals. Samples for physico-chemical attributes of the water were measured using a YSI multiparameter digital water quality meter, which took measurements of dissolved oxygen (%), pH, specific conductivity (μS/cm), and temperature (°C) averaged from depths 0.5 m and 1.5 m, as well as corrected calculations of dissolved oxygen (mg O_2_ L^-1^).

To track the effects of browning and eutrophication on water chemistry, 1-m deep water samples were collected from each mesocosm and from Lac Hertel. Vertical YSI probe profiles revealed stable water chemistry throughout each mesocosm (3 m depth), supporting the use of single-depth sampling for characterizing water column conditions (data not shown). Quantification of TP, TN and DOC concentrations were performed at the Groupe de Recherche Interuniversitaire en Limnologie (GRIL) laboratories based at Université du Québec à Montréal (see supplemental materials for analytical details). To evaluate the bioavailability (lability) of leachate-derived DOM relative to the naturally occurring DOM pool in Lake Hertel, we measured DOC biodegradation through a separate set of incubations starting on the first week of the experiment (August 31^st^) (see supplemental materials for analytical details).

Lastly, eDNA samples were collected at weeks 1, 5, and 9 to assess the composition of zooplankton, phytoplankton and bacterial communities (using primer sets targeting the COI, 18S rRNA and 16S rRNA genes, respectively). Grab samples of 1 L of water were collected from each mesocosm enclosure using a Nalgene amber, wide mouth bottle, prewashed overnight in an acid bath (HCl 10%), and wearing clean gloves. Water samples were then filtered through a 120 μm mesh and passed through a 0.22 μm 47 mm diameter filter using a manifold setup with a vacuum pump (all filtration apparatus was prewashed with 20% household bleach and rinsed three times with Milli-Q water). Water was filtered until the 0.22 μm filters clogged. The filters were then cut in half, with each half folded into a 2 mL cryovial and placed on dry ice in the field lab. Samples were transported the same day and stored at -80°C at the Environment and Climate Change Canada (ECCC) facilities in Montreal, Quebec. Filters were kept at -80°C until DNA extraction. Details on DNA extraction, amplicon sequencing and bioinformatics are described in the Supplementary Information.

*Statistical analyses.* All statistical analyses were conducted in R version 4.2.3 (R Development Core Team, 2023). To first explore temporal turnover in communities, for each trophic level (bacterioplankton, phytoplankton, and zooplankton), we calculated the Local Contribution to Beta Diversity (LCBD) using the *beta.div* function from the {adespatial} package (Dray et al. 2025). LCBD values are mathematically identical whether calculated on amplicon sequence variants (ASVs), genus- or other taxonomic-level data, as they reflect site-level contributions to compositional turnover. LCBD values were plotted against sampling time (categorical: T1, T5, T9) to determine if and when community composition deviated from average (centroid). To then explore whether browning (DOC) and eutrophication (TP+TN) treatments altered community composition, we performed a permutational multivariate ANOVA (PERMANOVA) (using the *adonis2* function in the {vegan} package; Oksanen et al. 2025) on Hellinger-transformed abundance matrices for each trophic group, with treatments as main factors. A Principle Coordinates Analysis (PCoA) was used to visualize the PERMANOVA results using the *cmdscale* function in Base R. To then highlight the taxa most responsive to treatments within each trophic level, we applied a Principal Response Curve (PRC) (using the *prc* function of the {vegan} package; Oksanen et al. 2025), which is a multivariate redundancy analysis (RDA) constrained by treatment and conditional on time. The first RDA axis scores (shown in the PRC figure output) indicate taxa showing clear increases or decreases in response to treatments, as well as the timing of peak responses and the treatment levels associated with the greatest effects. PERMANOVA and PRC analyses were conducted at multiple taxonomic levels for each community (genus to phylum). Exploratory, post-hoc plots of the relative abundance of dominant taxa (those with largest RDA axis 1 scores from the PRC) versus treatment concentrations were generated to illustrate potential thresholds. We therefore applied a changepoint analysis (using the *chngptm* function of the {chngpt} package; Fong et al. 2017) to interpret threshold responses in a manner consistent with our experimental design. Finally, DOC lability was analyzed using an ANCOVA on DOC concentrations as a function of incubation period (days) for each treatment combination, to verify comparability of DOC lability across treatments.

## Results

### Experimental treatments and environmental conditions

The addition of leaf litter and topsoil leachates enriched DOC in mesocosms to reach our target intermediate and high DOC levels. Our biodegradation incubations further indicated that the pool of DOC was similarly bioavailable between control and DOM-enriched mesocosms (Figure S1). Leachates were naturally rich in nutrients, leading to TP+TN spikes in the DOM-enriched mesocosms, reflecting the realistic co-enrichment associated with browning.

Consequently, starting from a mesotrophic control (lake water; mean TP = 13.1 ± 1.9 µg L^-1^), the *high DOC – no nutrient* addition combination was initially within the eutrophic range (mean TP = 79.6 µg L^-1^ ± 1.7 µg L^-1^), while hypereutrophic conditions (mean TP = 154.86 ± 8.44 µg L^-1^) were reached in the *high DOC – high nutrient* treatment (Figures 2, S2). Over the course of the experiment, both DOC and TP+TN concentrations declined, with treatments maintaining their rank order (Figure S3). By week 3, concentrations became more similar across treatments (Figure 2), suggesting that mesocosm mixing (i.e. no thermal stratification; data not shown) allowed natural attenuation processes (including nutrient uptake by organisms, microbial decomposition, photodegradation, and physical processes) to proceed uniformly across all treatments.

**Figure 2.**
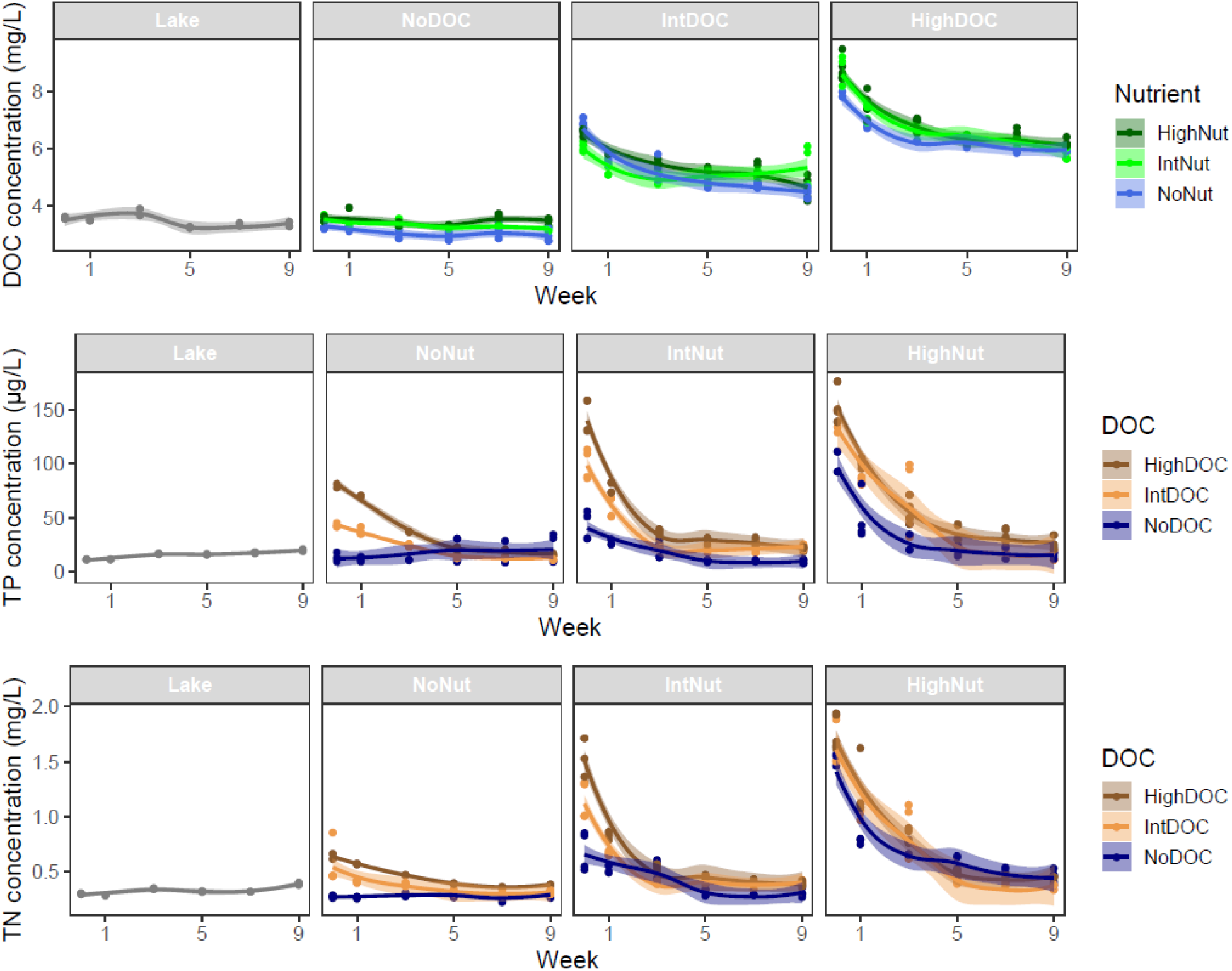
Time series **of a)** Dissolved organic carbon (mg L^-1^), **b)** Total phosphorus (µg L^-1^), and **c)** total nitrogen (mg N L^-1^) concentrations per browning and eutrophication treatments over the nine-week experiment.

### Community turnover

Community turnover, reported as the Local Contribution to Beta Diversity (LCBD), varied among trophic groups, with some slight differences in temporal patterns (Figure S4a-c). For zooplankton, the contribution of sites to community turnover (LCBD values) showed marginal variation among sampling dates (F = 2.99, p = 0.059), suggesting moderate temporal restructuring in community composition (Figure S4a). Phytoplankton turnover was more variable within sampling dates than among, and the change through time was not statistically significant (F = 0.973, p = 0.385; Figure S4b). For bacteria, beta diversity likewise did not differ significantly over time (F = 0.93, p = 0.401; Figure S4c), and as noted with phytoplankton, variability was greater within than among sampling dates.

### Effect of treatments on community composition

Taxonomic resolution was adjusted among organismal groups to account for differences in taxonomic richness. Because bacterial communities are highly speciose, analyses were conducted at the phylum level. Phytoplankton, which remain relatively speciose, were analyzed at the order level, whereas zooplankton were examined at the finest achievable taxonomic resolution due to their lower species richness. Exploratory analyses performed at alternative taxonomic resolutions yielded comparable results and did not influence the overall conclusions.

PERMANOVA indicated that browning (DOC) and eutrophication (TP+TN) treatments influenced community composition, with some slight differences among trophic groups. Zooplankton assemblages were significantly but modestly affected by DOC additions (F = 2.26, R² = 0.076, p = 0.019), whereas the effect of nutrients was marginal (F = 1.67, R² = 0.056, p = 0.094; Figure 3a). The DOC × nutrient interaction was likewise marginal (F = 1.67, R² = 0.112, p = 0.062). Pairwise comparison showed that communities in the DOC controls (lake water) differed significantly from the intermediate and high DOC additions (p < 0.05).

**Figure 3.**
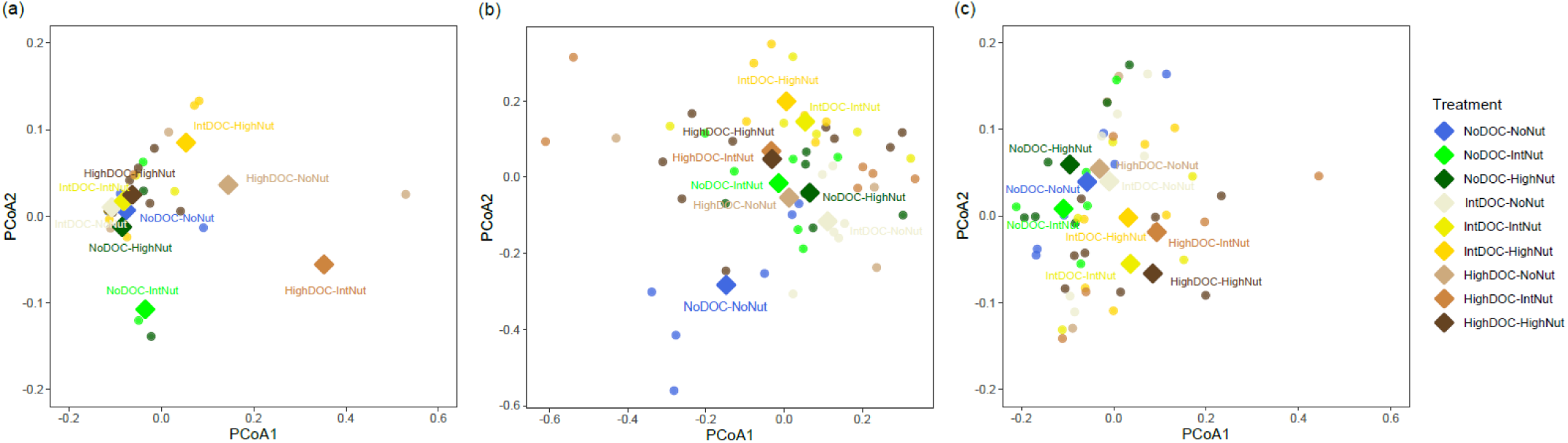
Principal coordinates analysis (PCoA) of mesocosm communities at different trophic levels: **a)** genus-level zooplankton (DOC +, Nut ∼, DOCxNut ∼), **b)** order-level phytoplankton (DOC +, Nut +, DOCxNut -), and **c)** phylum-level bacterioplankton (DOC +, Nut +, DOCxNut -). Ordinations illustrate community composition patterns corresponding to PERMANOVA results. Each point represents a mesocosm, colored by DOC × nutrient treatment, with centroids indicated by diamonds. (+; significant effect, -; no effect, ∼; marginal effect)

For phytoplankton, both DOC and nutrients significantly shaped community composition (DOC: F = 2.97, R^2^ = 0.10, p = 0.001; nutrients: F = 2.78, R^2^ = 0.09, p = 0.006). However, no significant interactive effects were detected between treatments (F = 0.95, R^2^ = 0.07, p = 0.523). Pairwise comparisons indicated that communities in the controls (mesotrophic lake water with no treatment additions) differed from all enriched treatments, whereas intermediate and high addition treatments did not significantly differ from each other (Figure 3b). Thus, enrichment relative to baseline drove community shifts, but increasing beyond intermediate levels had little additional effect on phytoplankton community composition.

For bacterioplankton, DOC had the strongest effect on community composition (F = 3.42, R² = 0.12, p = 0.0005), again distinguishing the no DOC controls from other levels (Figure 3c). Nutrient addition treatments had a more selective effect (F = 1.84, R² = 0.06, p = 0.045), distinguishing only between the extreme contrasts (no vs high nutrient additions). The interaction between DOC and nutrients was once again non-significant (F = 0.84, R² = 0.06, p = 0.67).

Overall, the PERMANOVA showed that all communities responded to treatments, with DOC being a primary driver of zooplankton community composition, and both DOC and nutrients contributing to shifts in phytoplankton and bacterial communities.

### Browning and eutrophication threshold responses

The PRC ordination constrained by treatment, conditional on time, showed that DOC consistently drove the strongest shifts in plankton and bacterial composition, with nutrient enrichment (TP+TN) modulating responses, and thresholds generally detected at intermediate treatment levels (Figure 4).

**Figure 4.**
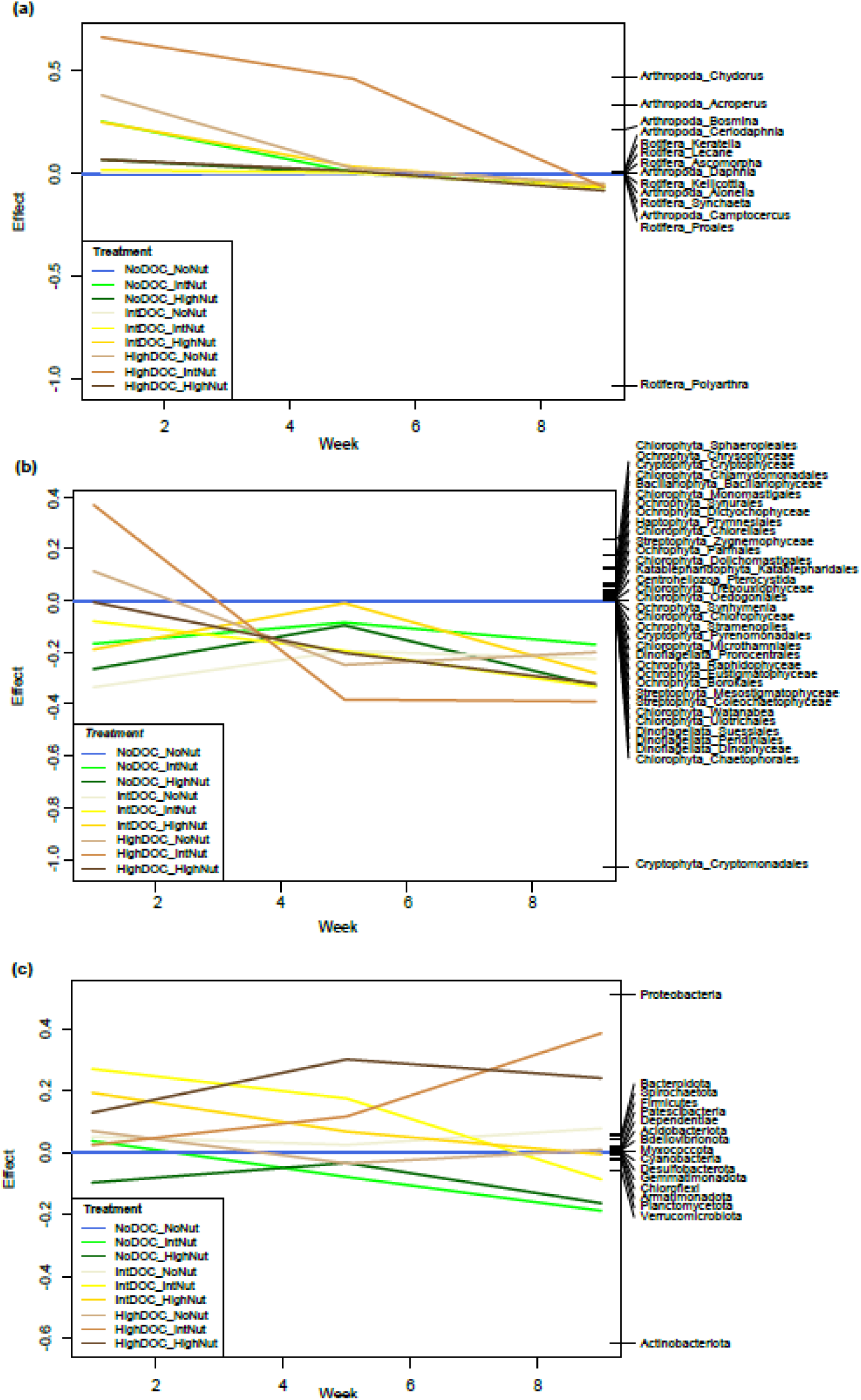
Principal Response Curves (PRC) showing the effect of browning × eutrophication treatments on community composition over time, conditional on week, for (a) zooplankton genera, (b) phytoplankton orders, and (c) bacterial phyla. Curves are colored by treatment level, illustrating how each community composition responses diverge from the control over the course of the experiment.

Zooplankton community composition experienced a pronounced shift among treatments at the start of the experiment (one week post treatment additions); however, by week 9 all treatments converged to the lake-water control (coefficients near zero or slightly negative; Figure 4a). At the start of the experiment, Cladocerans (*Bosmina*, *Chydorus*, *Acroperus*, to some extent *Ceriodaphnia*) peaked under conditions of intermediate nutrients and high DOC, while rotifers (especially *Polyarthra*) were more strongly associated with high nutrients and high DOC spikes. Changepoint analyses further showed that, among the taxa with the largest PRC loadings (i.e. showing the most changes in abundance across treatments), breakpoints in relative abundances were observed in the range of 6.5 to 7.0 mg L^-1^, with a median of 6.5 mg L^-1^ (Figures S4d, 5a). The median breakpoints for TP for the same taxa were noted at 69 ug L^-1^ (not shown).

The PRC analysis applied to phytoplankton data showed that DOC treatments had the largest effect, followed by nutrients (Figures 4b). At the start of the experiment, dominant positive responders included Chrysophyceae, Cryptophyceae, Chlamydomonadales (mixotrophic), and Sphaeropleales (green algae involved in nutrient uptake). In contrast, Cryptomonadales (typically adapted to light limitation but sensitive to nutrient competition) showed the lower PRC scores. By week 5 onwards, however, we noted a gain in Cryptomonadales within the high DOC treatments (treatment lines shift to lower quadrant). This pattern aligned with field observations: Cryptomonadales exhibited a unimodal response to additions, with low relative abundances under high DOC and nutrients conditions at the start of the experiment, but recovered as DOC and nutrient concentrations reached intermediate levels (Figure 5b). Changepoint analysis on the taxa with the strongest loadings showed similar DOC threshold as zooplankton (median 6.5 mg L^-1^: range 3.3 to 7 mg L⁻¹; Figure S4e, 5b). The median TP breakpoint was noted at 62 ug L^-1^ (not shown).

**Figure 5.**
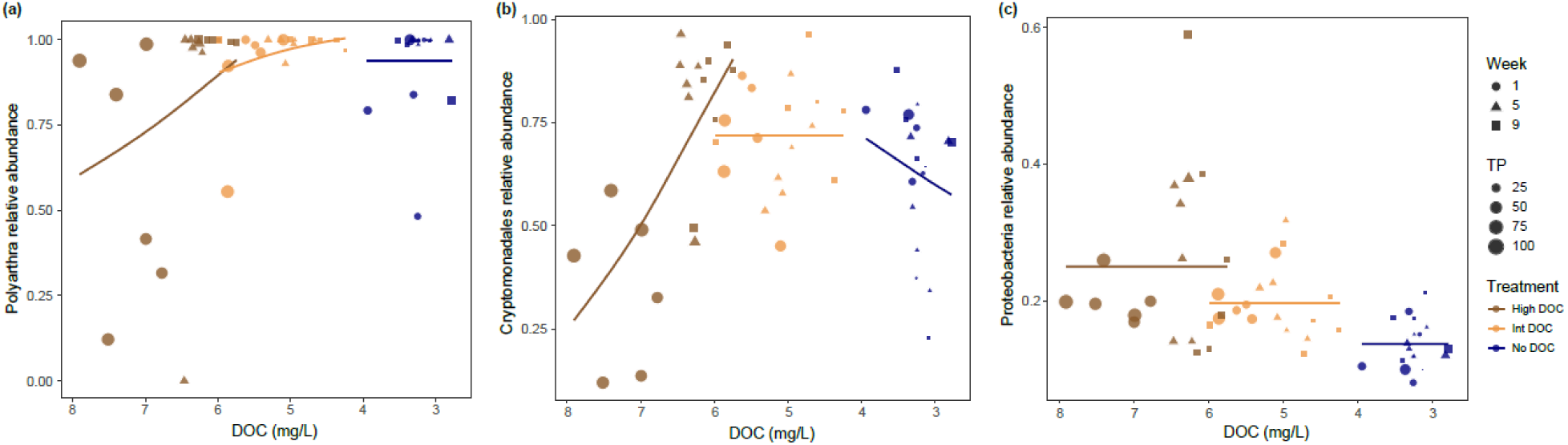
Relative abundances of *Polyarthra* (zooplankton, a), Cryptomonadales (phytoplankton, b), and Proteobacteria (bacterioplankton, c) versus DOC concentrations. Point size represents total phosphorus (TP) concentration, point shape indicates sampling week, and color denotes DOC treatment level. Fitted lines show the relationship between relative abundance and DOC for each DOC treatment. The x-axis displays decreasing DOC concentrations, representing the timeline of the pulse effect.

Finally, the treatment-time constrained ordination of bacterial communities identified an effect of DOC, though changes in relative abundance were less pronounced (Figures 4c). Positive responders included Proteobacteria and Firmicutes, while Actinobacteriota declined. Breakpoints for dominant bacterial phyla ranged from 3.1 mg L⁻¹ (Verrucomicrobiota) to 6.5 mg L⁻¹ (Bacteroidota), spanning a broader range than plankton and yielding a slightly lower median (4.9 mg L⁻¹; Figure S4f). Temporally, bacterial shifts first occurred at the start of the experiment (week 1) under intermediate DOC and nutrients, followed by later shifts in weeks 5 to 9 that coincided with a pronounced drop in DOC and nutrient concentrations in the high DOC treatments (Figure 2) and with the formation of periphyton on the mesocosm walls (observed on week 5 and onwards). The median TP thresholds for the phyla with the largest loadings were observed at 31 ug L^-1^ on average (ranging from 17 to 62 ug L^-1^), thus lower (later) than the plankton groups (not shown), highlighting a gradient of nutrient sensitivity across trophic groups.

Taken together, as DOC declined after the initial pulse, plankton responded rapidly, with rotifers replacing cladocerans, and cryptophytes replacing chlorophytes on week 5 onwards (Figures S4d-e, 5a-b). In contrast, bacterial changes were less synchronous: we noted multiple shifts with responses variable across phyla, and lacked the sharp patterns observed for plankton (Figures S4f, 5c). To assess whether the muted bacterial response resulted from the coarser taxonomic resolution used for this group, the PRC at order and phylum level were compared. Results highlighted the same relative abundance patterns (not shown), indicating that the weaker bacterial signal is not an artifact of taxonomic resolution.

## Discussion

Browning and eutrophication are recognized drivers of freshwater planktonic community change; yet more integrative research is needed to clarify their joint impacts on community threshold responses and cross-trophic dynamics. Disentangling these effects may require experimental approaches that capture both pulse-driven (i.e. immediate) and sequential (i.e. long-lasting) multitrophic responses to individual and combined increases in lake DOC and nutrients (Orr et al. 2020). In this respect, in-lake mesocosm experiments provide a controlled, ecologically realistic approach to isolate the relative contributions of drivers, identify responsive taxa, and characterize community responses across the range of DOC and nutrient concentrations achieved (Stewart et al. 2013). We addressed this by using eDNA amplicon sequencing to assess how pulse treatments of browning (range of achieved DOC concentrations: 2.7-9.5 mg L^-1^) and eutrophication (range of achieved nutrient concentrations: 7.4-176.4 μg P L^-1^ and 0.23-1.9 mg N L^-1^) led to changes in bacterioplankton, phytoplankton and zooplankton community composition over nine weeks (late July-October). Our study revealed that DOC was a strong driver of community change across all trophic levels, with nutrient modulating responses from bacterial and algal communities. Thresholds generally occurred under intermediate treatments, where responsive groups included mainly proteobacteria, cryptophytes and rotifers.

### Beta diversity across trophic groups

We calculated LCBD to quantify how much each sample contributed to overall compositional variance. For zooplankton, LCBD peaked in week 1 following DOC and nutrient pulses, then declined and remained low through week 9, suggesting early restructuring (Figure S4). Phytoplankton and bacterioplankton showed consistently high within-week variability with no clear temporal effect. These patterns supported experiment-wide analyses rather than weekly comparisons and justified subsequent PERMANOVAs and constrained ordinations to identify species-specific responses to treatments. Past studies have reported either a decrease in temporal beta-diversity with eutrophication (e.g. for macroinvertebrates; Cook et al. 2018) or nonmonotonic responses to lake browning (e.g. for bacterial assemblages; Fontaine et al. 2021). Here, however, we found that the response to treatments, irrespective of time was more important.

### Response threshold concentrations and sensitive taxa across the food web

PERMANOVA indicated that experimental treatments influenced community composition across trophic levels, though to different degrees. Zooplankton were primarily affected by DOC, with nutrients having a marginal effect. In contrast, phytoplankton responded to both DOC and nutrients, but with little additional change beyond intermediate treatments. This pattern suggests that phytoplankton may respond more strongly to shifts in the form of basal resources than to further increases in their overall quantity (Reinl et al. 2022). Bacterial communities were strongly influenced by DOC, whereas responses to nutrients were limited to extreme enrichment conditions (difference observed between no vs high nutrient addition treatments only). Overall, these patterns indicate that DOC was the dominant driver of community shifts, whereas nutrient effects varied between trophic levels.

To identify which taxa drove patterns of community turnover and how they responded to DOC and nutrient pulses over time, we then applied a Principal Response Curve (PRC) analysis. This treatment-by-time constrained ordination identified taxon-specific responses, notably to DOC enrichment. By the end of the first week (day 7) following the initial treatment pulse (day 1), Cladocera (*Chydorus*, *Acroperus*, and *Bosmina*) were associated with intermediate nutrient combined with high DOC treatments, while rotifer zooplankton (*Polyarthra*) were associated with the high nutrient-high DOC combination. This pattern is consistent with the addition of terrestrially derived organic matter (i.e. allochthonous carbon) favouring the microbial-detrital food chain and thereby favouring small detritivorous and bacteriovorous microzooplankton (such as rotifers) (Berggren et al. 2014). In line with this interpretation, bacterial groups associated with carbon cycling (Proteobacteria), anaerobic degradation of complex organic matter (Firmicutes), and biofilm formation (Bacteroidota) reached peak abundance under intermediate to high DOC treatments during week 1. This pattern suggests that elevated DOC promoted anaerobic microsites within biofilms and on suspended particles. In contrast, phyla often considered oligotrophic, thriving in nutrient-poor environments (Actinobacteriota), involved in the degradation of plant-derived organic matter (Verrumicrobiota), or thriving in nitrogen-rich environments (Planctomycetota) (Zhang et al. 2024; Rakitin et al. 2024) were associated with the no-DOC combined with intermediate to high nutrient treatments. By week 5, as DOC declined below ∼5 mg L⁻¹, mesocosms with sufficient nutrients continued to support rotifers and the dominance of small phytoplankton such as cryptophytes (Cryptophyceae, Cryptomonadales), which are commonly consumed by rotifers (Liang et al. 2022).

Overall, the community compositional responses are consistent with a temporal shift in zooplankton resources, with elevated DOM and nutrients first fueling the microbial-detrital food chain (brown pathway) and then later phytoplankton (green pathway). This broadly supports our initial hypotheses and aligns with predictions by Creed et al. (2018), who suggested that microbial biomass would increase under elevated DOC, whereas phytoplankton would dominate at lower DOC concentrations. Our results also echo prior observations that rotifers tend to benefit more when the microbial loop is active and bacteria constitute a significant part of available food (Berggren et al. 2014).

The DOC thresholds leading to the greatest changes in dominance patterns further highlight differences among DOC-treated communities and their potential reliance on bacterial versus phytoplankton loops. Notably, community responses to DOC peaked at 6.5 mg L⁻¹ within plankton and later at 4.9 mg L⁻¹ within bacteria (where DOC concentrations at or below 5 mg L⁻¹ were recorded in either the no-DOC treatments or the intermediate DOC–no nutrient treatments from week 5 onwards).

Given that DOC decreased over time, the sequence of thresholds is consistent with a bottom-up process, where shifts in resource availability (DOC and nutrients) drove sequential changes in microbial, phytoplankton, and zooplankton communities: high DOC initially triggered a microbial-loop response (rotifers favoured, cladocerans suppressed), followed by a rise in small phytoplankton that supported cladoceran populations, and later bacterial responses as DOC declined and temperatures dropped towards season’s end. Notably, bacterioplankton composition appeared to be driven primarily by treatment effects rather than temporal dynamics; individual taxa (e.g., Proteobacteria) showed consistent relative abundances across the DOC gradient within each treatment, with little change over time. The bacterial thresholds observed herein are consistent with those reported in a eutrophic mesocosm experiment, where 4.6 mg L⁻¹ DOC was associated with an abrupt change in cyanobacteria biomass (Feuchtmayr et al., 2019). Although Xie et al. (2020) also attributed bacterial community changes to low temperature and dissolved oxygen, we found, in a post-hoc analysis, only a marginal effect of temperature on bacteria (Figure S5). However, when considering only the high DOC treatment, temperature significantly influenced Actinobacteriota (F = 5.03, p = 0.04), with relative abundances dropping by as much as 40%. This pattern may reflect an interaction between DOC and temperature, where the metabolic activity of heterotrophic bacteria (like Actinobacteriota) declined as organic carbon and temperatures dropped, while other bacteria (facultative anaerobes, or taxa favored by cooler, oxygenated water) remained stable. Observed thresholds across trophic groups may also reflect additional factors, including changes in light availability limiting primary production (Seekell et al., 2015) and variation in food quality (e.g., macro- and micronutrients). Browning in unproductive lakes more strongly affects heterotrophic and mixotrophic zooplankton than in eutrophic lakes, primarily due to greater reliance on terrestrial carbon subsidies in low-nutrient systems (Carpenter et al., 2005; Kelly et al., 2016; Carpenter et al., 2022). Overall, the dynamics observed over the course of the nine-week experiment highlight that while DOC has the potential to drive strong compositional shifts, community responses are also shaped by interactions with nutrient availability and physico-chemistry, emphasizing the need for holistic analyses that consider multiple co-occurring drivers when scaling up to natural lake ecosystems across the landscape.

## Future directions

This study demonstrates the complex interactions between browning and eutrophication in shaping freshwater communities, identifying treatment thresholds that drove shifts in bacterioplankton, phytoplankton, and zooplankton community assemblages. While short term mesocosms provide valuable mechanistic insights, they are simplified systems and may not capture all natural interactions or seasonal variability. Additionally, the two-month duration of the experiment was likely insufficient to capture adaptive or ecological processes that occur over longer timescales in natural lake ecosystems. Building on these results, subsequent work should integrate *in situ* lake monitoring to validate thresholds and multi-trophic responses across diverse lakes and seasons. Further research could also examine how browning and eutrophication influence contaminant dynamics, such as mercury and cyanotoxin bioaccumulation, mediated by DOC inputs and algal blooms, thereby linking community shifts to ecosystem health and water quality.

## Acknowledgements

This study is a contribution to the Climate Change and Atmospheric Pollution (CCAP) research program of Environment and Climate Change Canada (ECCC). The authors would like to thank the personnel at 1) ECCC: Conrad Beauvais, Suzanne Couture, Laurie Mercier and Benoit Fortin, 2) McGill University: Paul MacKeigan, David Zilkey, Jennifer Pham, Michelle Cheng and Egor Katkov, and 3) other universities: Cindy Paquette (UQAM) and Sufyan Mirza (Université de Sherbrooke). We thank Katherine Velghe (GRIL; UQAM) for her help and support on nutrient analyses. We also thank the Gault Nature Reserve for supporting us in providing the facilities and equipment to conduct this experiment on the mesocosms dock at Lac Hertel. Finally, we thank the Canada Research Chair program for supporting IGE and NSERC for supporting M-ÈM and M-PH.

## Data statement

Data available on request from the authors.

## Author Contribution Statement

ML led the field sampling, data acquisition, statistical analyses, and drafting of the manuscript. ZET led the study’s conception and co-led statistical analyses and drafting of the manuscript. IGE contributed to the study’s conception, data interpretation and drafting of the manuscript. SK contributed to bioinformatics, data interpretation (eDNA) and editing the manuscript. MA contributed to data interpretation and editing of the manuscript. M-ÈM contributed to the study’s conception, data interpretation (eDNA), and editing the manuscript. M-PH contributed to the study’s conception, data interpretation (DOC, zooplankton), and editing the manuscript. All authors approved the final submitted manuscript.

## Abbreviations

DOC: Dissolved Organic Carbon
TP: Total Phosphorus
eDNA: environmental
DNA, PRC: Principal Response Curve
LCBD: Local Contribution to Beta Diversity
PERMANOVA: Permutational Multivariate Analysis of Variance

