## Supplemental Material for "Threshold responses of multi-trophic freshwater communities to browning and eutrophication"

**Supplemental Information for methodology:**

***Experimental design***

On day zero of the experiment, for the intermediate eutrophication treatment we added 4.289 g of $NaNO_3$, 0.104 g of $KH_2PO_4$, and 0.133 g of $K_2HPO_4$ and for the high treatment added 8.578 g of $NaNO_3$, 0.207 g of $KH_2PO_4$, and 0.265 grams of $K_2HPO_4$.

For the browning treatments, we added our leaf and soil brew to dedicated enclosures. Prior to adding leachates, we first removed an equal volume of water from the treated enclosures, filtering the removed water through a 120 µm mesh and then placed back the particulate material into the mesocosm so as to maintain the plankton community of interest.

***Sample processing***

TP and TN samples were oxidized (autoclave, at 121$^{o}$C) with a potassium persulfate digestion. TP was measured via colorimetric detection using the molybdenum method at 890 nm with a spectrophotometer (Ultrospec 2100 pro, Biochrom). TN concentrations were determined using a continuous flow analyzer coupled with a cadmium reactor (OI Analytical, Flow Solution 3100). DOC concentrations (with water samples pre-filtered at 0.45µm) were quantified with an automated TIC-TOC Analyzer (Aurora 1030, OI Analytical) using persulfate oxidation.

***Dissolved Organic Carbon (DOC) bioavailability (lability) test; for leachate-derived DOM used as treatment to increase DOC in mesocosms***

In short, we filtered water samples through pre-burned Whatman GF/D 47mm filters, then stored in the dark at room temperature (20$^{o}$C) in pre-conditioned (acid-washed) glass amber bottles, with semi-screwed caps to allow for air circulation. We then tracked DOC biodegradation over a total of 37 days, collecting sub-samples every 3-4 days, acidified with sulfuric acid 5N and stored at 4°C until analysis at the GRIL laboratories for DOC concentrations (Hébert et al. 2022).

***DNA extraction and High-throughput sequencing (HTS).***

DNA was extracted (DNeasy PowerSoil DNA isolation kit) and sequenced (Illumina MiSeq PE 300 bp) at Genome Quebec facilities (Montreal, Canada). The bacterial community was targeted by amplifying a 291 bp fragment of the V4 region of the 16S rRNA gene using the universal bacterial primers U515F primer (5′- GTGYCAGCMGCCGCGGTAA-3′) and E786R (5′-GGACTACNVGGGTWTCTAAT-3′) (Caporaso et al., 2011). For the phytoplankton community, a 478 bp fragment of the 18S rRNA gene V7 region was amplified using the universal microeukaryotes primers 960F (5′-AGTGGCGGACGGGTGAGTAA-3′) (Gast et al., 2004) and NSR1438 (5′-GCTGCTGGCACCAGACTTGC-3′) (Hadziavdic et al., 2014). Finally, for the zooplankton (metazoa) community, a 313 bp fragment of the mitochondrial Cytochrome c Oxidase subunit I gene (COI) was amplified using the universal metazoan primers mlCOIintF (5′-GGWACWGGWTGAACWGTWTAYCCYCC-3′) (Leray et al., 2013) and jgHCO2198 (5′-TAIACYTCIGGRTGICCRAARAAYCA-3′) (Geller et al., 2013).

***Bioinformatics.***

Raw sequencing reads were first processed using Cutadapt version 2.6 (Martin, 2011) to remove primers and adapters. To recover amplicon sequence variants (ASVs), the DADA2 pipeline (Callahan et al., 2016) was then applied in R (version 4.2.3; R Development Core Team, 2023). This involved trimming and filtering forward and reverse reads, dereplicating, removing chimeras, and finally merging paired ends. Taxonomic assignment for 16S rRNA was performed against the SILVA database (Glöckner et al., 2017). The 18s rRNA taxonomic assignment was realized using the PR2 database (Guillou et al., 2013). Finally, COI taxonomic assignment was performed with both the RDP (Ribosomal Database Project) classifier (Wang et al., 2007) and the MIDORI database (Leray et al., 2018). COI pseudogenes were removed by using MACSE (Ranwez et al., 2018) to align all putative COI genes against representative alignments of MIDORI COI vertebrate and non-vertebrate sequences, and excluding any sequences with stop codons, frameshifts or more than five deletions (John Pearman, personal communication). For the zooplankton community, only the Phyla Rotifera, Cnidaria, Annelida, Mollusca and Arthropoda were retained from the COI dataset. For the cyanobacteria community, only the Cyanobacteria Phylum (n=143 taxa) was selected from the 16S dataset. For the microbial community, a total of 891 taxa were selected from the 16S dataset. To reduce the dimensionality of the datasets, we only retained taxa with a minimum occurrence of two observations and minimum relative abundance of 0.01% (using the *dropspc* function of the {labdsv} package; Roberts, 2023). All species matrices were Hellinger transformed to maintain a Euclidean distance matrix when performing multivariate statistical analyses.

**Supplemental Information: Research gaps and caveats**

*DOM composition.* In this study, DOM constituents were not analyzed optically or otherwise; therefore, we have no robust indication of how DOC composition may have changed over the nine-week period. Nonetheless, we conducted lability incubations in parallel to our main experiment to measure and compare DOC biodegradability among treatments. Overall, we determined that the bioavailability of DOC in leaf litter and topsoil leachates was similar among treatments, although it remains unknown if this carbon was incorporated into bacterial biomass or lost through respiratory fluxes. Future studies should aim to better characterize the main forms of DOC, as this would reveal its origin, aromaticity, and molecular weight, ultimately providing a more accurate estimation of DOM's faith and binding capacity (Findlay, 2003; Castan et al., 2020).

**Supplemental figures for results:**

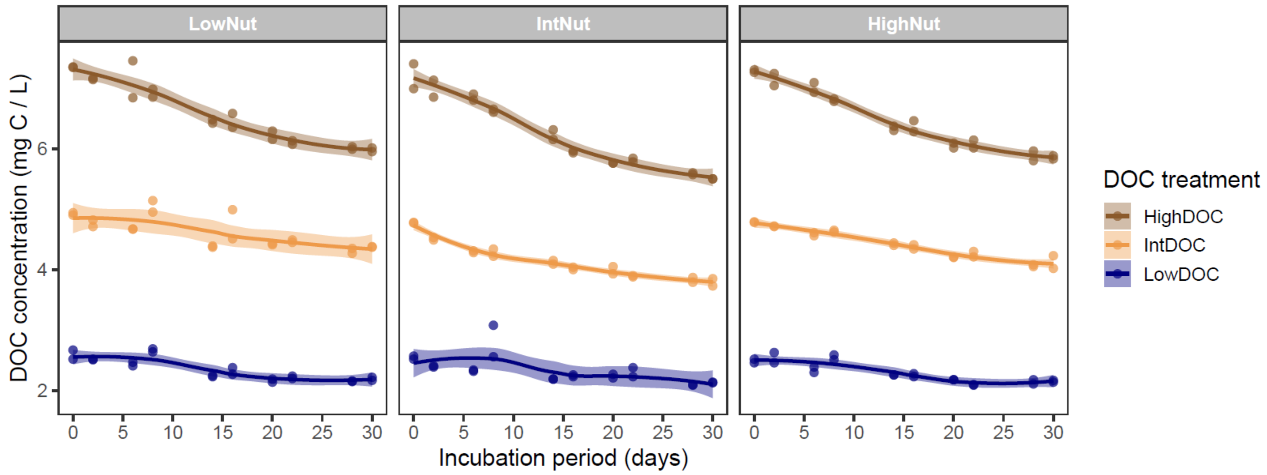


**Figure S1.** Dissolved organic carbon biodegradation over time for each eutrophication and browning treatment levels.

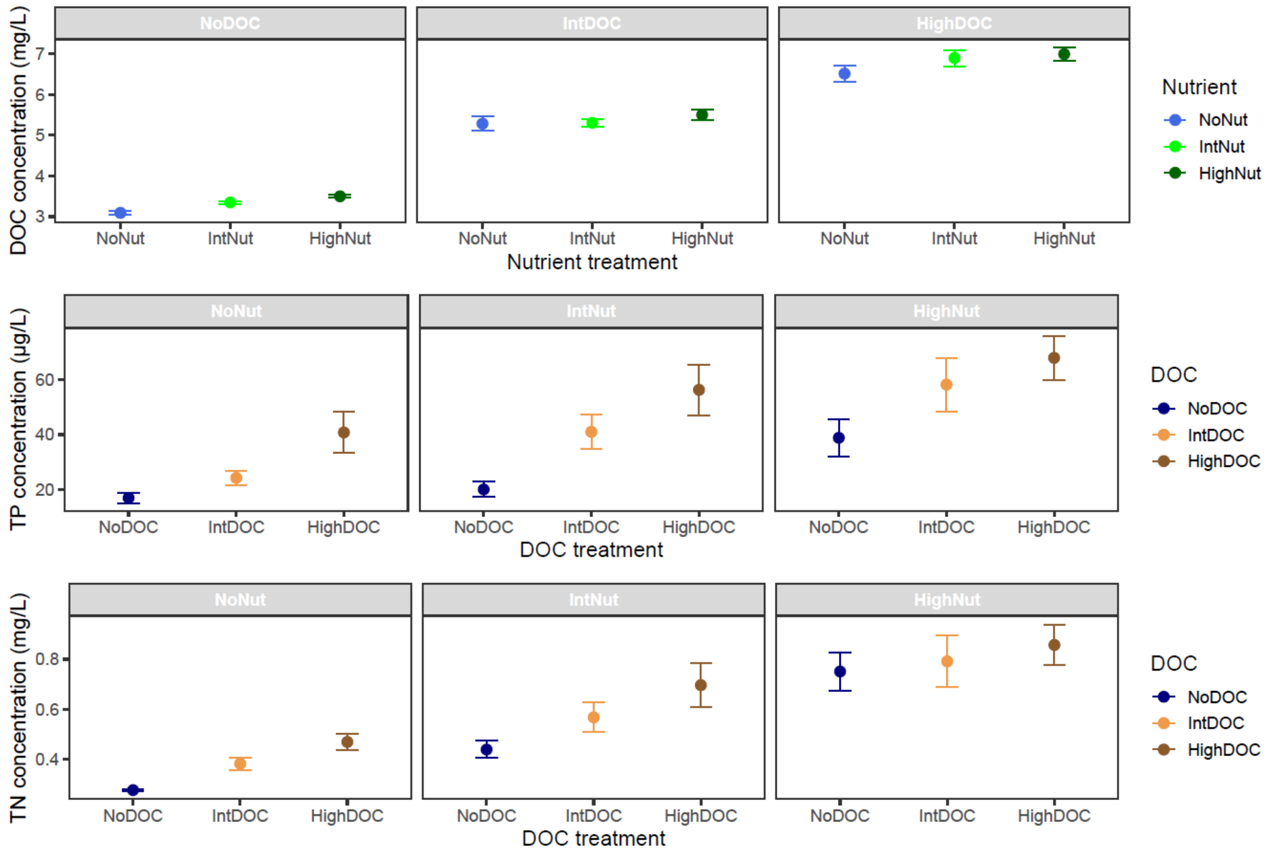


**Figure S2.** Mean ± standard error of **a)** dissolved organic carbon (DOC; mg $L^{-1}$), **b)** total phosphorus (TP; µg $L^{-1}$), and **c)** total nitrogen (TN; mg N $L^{-1}$) concentrations across browning and eutrophication treatments over the nine-week experiment.

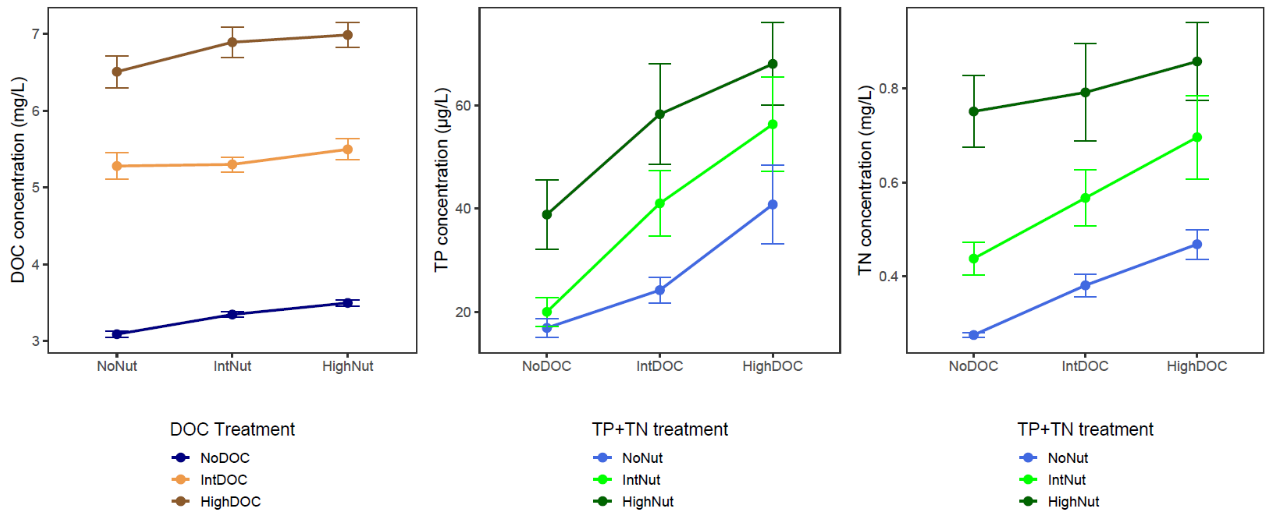


**Figure S3.** Mean ± standard error of **a)** dissolved organic carbon (DOC; mg $L^{-1}$) by nutrient treatments, and **b)** total phosphorus (TP; µg $L^{-1}$) or **c)** total nitrogen (TN; mg N $L^{-1}$) by DOC treatments, illustrating that within each treatment level, concentrations remained in consistent rank order.

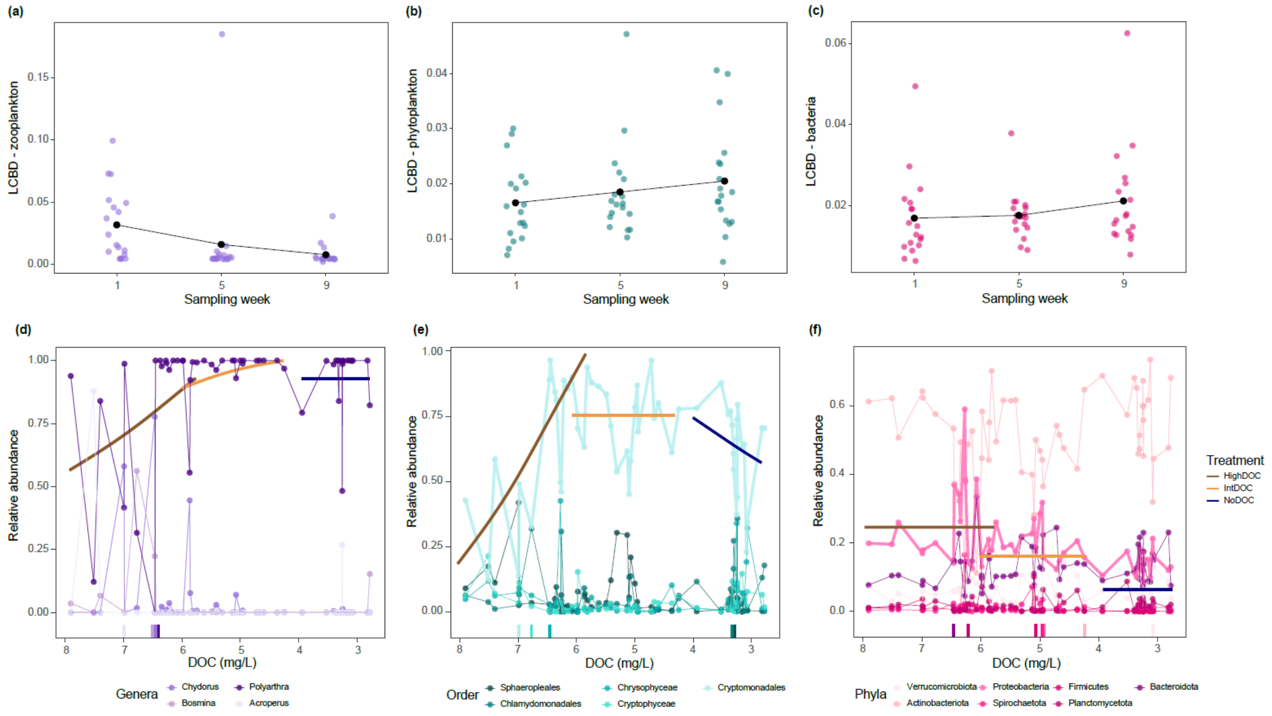


**Figure S4.** Community turnover (a-c) and abundance responses (d-f) across trophic groups. Local Contribution to Beta Diversity (LCBD) are shown for **a)** zooplankton, **b)** phytoplankton, and **c)** bacteria over time. Panels **(d–f)** show relative abundances (zooplankton genera in d, phytoplankton orders in e, dominant bacterial phyla in f) versus DOC for the taxa with highest Principal Response Curve (PCR) loadings (see Figure 4). Tick marks on the x-axis of d-f indicate changepoints detected for each taxon, and colored lines (blue, tan, brown) show linear fits of relative abundance versus DOC for the browning treatment levels (no, intermediate, high DOC), illustrated here for *Polyarthra* (zooplankton), Cryptomonadales (phytoplankton), and Proteobacteria (bacterioplankton) (see Figure 5).

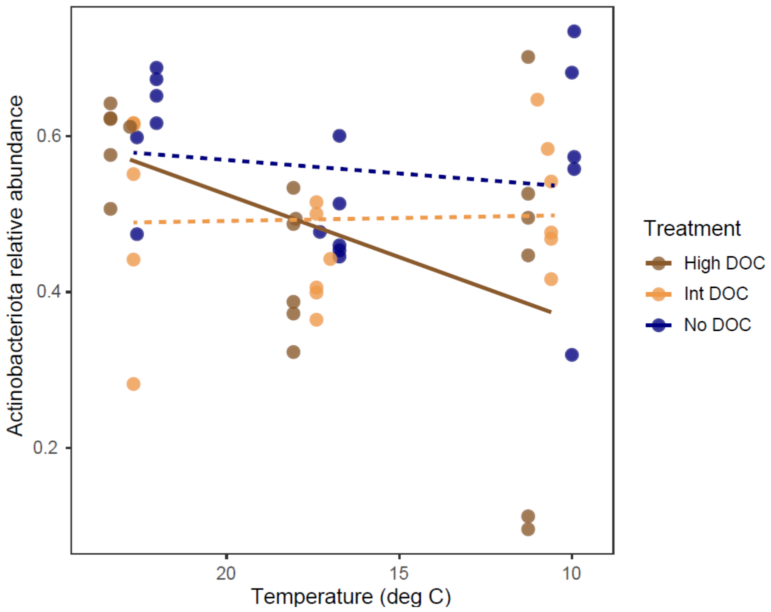


**Figure S5.** Relative abundance of Actinobacteria versus water temperature. Points are colored by DOC treatment, and linear regression lines show the relationship between relative abundance and temperature for each DOC treatment level. Dashed lines represent relationships that were not significant.

**Supplemental Information: References**

Bravo, A. G., Bouchet, S., Tolu, J., Björn, E., Mateos-Rivera, A., & Bertilsson, S. (2017). Molecular composition of organic matter controls methylmercury formation in boreal lakes. *Nature Communications*, *8*(1), 14255. https://doi.org/10.1038/ncomms14255

Callahan, B. J., McMurdie, P. J., Rosen, M. J., Han, A. W., Johnson, A. J. A., & Holmes, S. P. (2016). DADA2: High-resolution sample inference from Illumina amplicon data. *Nature Methods*, *13*(7), 581–583. https://doi.org/10.1038/nmeth.3869

Caporaso, J. G., Lauber, C. L., Walters, W. A., Berg-Lyons, D., Lozupone, C. A., Turnbaugh, P. J., et al. (2011). Global patterns of 16S rRNA diversity at a depth of millions of sequences per sample. *Proceedings of the National Academy of Sciences of the United States of America*, *108 Suppl 1*(Suppl 1), 4516–4522. https://doi.org/10.1073/pnas.1000080107

Castan, S., Sigmund, G., Hüffer, T., Tepe, N., Kammer, F. von der, Chefetz, B., & Hofmann, T. (2020). The importance of aromaticity to describe the interactions of organic matter with carbonaceous materials depends on molecular weight and sorbent geometry. *Environmental Science: Processes & Impacts*, *22*(9), 1888–1897. https://doi.org/10.1039/D0EM00267D

Findlay, S. (2003). Bacterial Response to Variation in Dissolved Organic Matter. In *Aquatic Ecosystems* (pp. 363–379). Elsevier. https://doi.org/10.1016/B978-012256371-3/50016-0

Gast, R. J., Dennett, M. R., & Caron, D. A. (2004). Characterization of Protistan Assemblages in the Ross Sea, Antarctica, by Denaturing Gradient Gel Electrophoresis. *APPL. ENVIRON. MICROBIOL.*, *70*.

Geller, J., Meyer, C., Parker, M., & Hawk, H. (2013). Redesign of PCR primers for mitochondrial cytochrome *c* oxidase subunit I for marine invertebrates and application in all-taxa biotic surveys. *Molecular Ecology Resources*, *13*(5), 851–861. https://doi.org/10.1111/1755-0998.12138

Glöckner, F. O., Yilmaz, P., Quast, C., Gerken, J., Beccati, A., Ciuprina, A., et al. (2017). 25 years of serving the community with ribosomal RNA gene reference databases and tools. *Journal of Biotechnology*, *261*, 169–176. https://doi.org/10.1016/j.jbiotec.2017.06.1198

Guillou, L., Bachar, D., Audic, S., Bass, D., Berney, C., Bittner, L., et al. (2013). The Protist Ribosomal Reference database (PR2): a catalog of unicellular eukaryote small sub-unit rRNA sequences with curated taxonomy. *Nucleic Acids Research*, *41*(Database issue), D597-604. https://doi.org/10.1093/nar/gks1160

Hadziavdic, K., Lekang, K., Lanzen, A., Jonassen, I., Thompson, E. M., & Troedsson, C. (2014). Characterization of the 18S rRNA Gene for Designing Universal Eukaryote Specific Primers. *PLoS ONE*, *9*(2), e87624. https://doi.org/10.1371/journal.pone.0087624

Hébert, M. P., Soued, C., Fussmann, G. F., & Beisner, B. E. (2022). Dissolved organic matter mediates the effects of warming and inorganic nutrients on a lake planktonic food web. *Limnology and Oceanography*, *68*, S23-S38. https://doi.org/10.1002/lno.12177

Houle, D., Khadra, M., Marty, C., & Couture, S. (2020). Influence of hydro-morphologic variables of forested catchments on the increase in DOC concentration in 36 temperate lakes of eastern Canada. *Science of The Total Environment*, *747*, 141539. https://doi.org/10.1016/j.scitotenv.2020.141539

Imtiazy, M. N., Paterson, A., Higgins, S., Yao, H., Houle, D., & Hudson, J. (2024). Has lake brownification ceased? Stabilization, re-browning, and other factors associated with dissolved organic matter trends in eastern Canadian lakes. *Water Research*, *269*, 122814. https://doi.org/10.1016/j.watres.2024.122814

Lázaro, W. L., Díez, S., Bravo, A. G., da Silva, C. J., Ignácio, Á. R. A., & Guimaraes, J. R. D. (2019). Cyanobacteria as regulators of methylmercury production in periphyton. *Science of The Total Environment*, *668*, 723–729. https://doi.org/10.1016/j.scitotenv.2019.02.233

Leclerc, M., Harrison, M. C., Storck, V., Planas, D., Amyot, M., & Walsh, D. A. (2021). Microbial Diversity and Mercury Methylation Activity in Periphytic Biofilms at a Run-of-River Hydroelectric Dam and Constructed Wetlands. *MSphere*, *6*(2), 10.1128/msphere.00021-21. https://doi.org/10.1128/msphere.00021-21

Leray, M., Yang, J. Y., Meyer, C. P., Mills, S. C., Agudelo, N., Ranwez, V., et al. (2013). A new versatile primer set targeting a short fragment of the mitochondrial COI region for metabarcoding metazoan diversity: application for characterizing coral reef fish gut contents. *Frontiers in Zoology*, *10*(1), 34. https://doi.org/10.1186/1742-9994-10-34

Leray, M., Ho, S.-L., Lin, I.-J., & Machida, R. J. (2018). MIDORI server: a webserver for taxonomic assignment of unknown metazoan mitochondrial-encoded sequences using a curated database. *Bioinformatics*, *34*(21), 3753–3754. https://doi.org/10.1093/bioinformatics/bty454

Li, Z., Chi, J., Shao, B., Wu, Z., He, W., Liu, Y., et al. (2022). Inhibition of methylmercury uptake by freshwater phytoplankton in presence of algae-derived organic matter. *Environmental Pollution*, *313*, 120111. https://doi.org/10.1016/j.envpol.2022.120111

Martin, M. (2011). Cutadapt removes adapter sequences from high-throughput sequencing reads. *EMBnet.Journal*, *17*(1), 10–12. https://doi.org/10.14806/ej.17.1.200

R Development Core Team. (2023). R: A language and environment for statistical computing. Retrieved from https://www.r-project.org/

Ranwez, V., Douzery, E. J. P., Cambon, C., Chantret, N., & Delsuc, F. (2018). MACSE v2: Toolkit for the Alignment of Coding Sequences Accounting for Frameshifts and Stop Codons. *Molecular Biology and Evolution*, *35*(10), 2582–2584. https://doi.org/10.1093/molbev/msy159

Roberts, D. W. (2023). labdsv: Ordination and Multivariate Analysis for Ecology (Version 2.1-0) [Data set]. https://doi.org/10.32614/CRAN.package.labdsv

Wang, Q., Garrity, G. M., Tiedje, J. M., & Cole, J. R. (2007). Naive Bayesian classifier for rapid assignment of rRNA sequences into the new bacterial taxonomy. *Applied and Environmental Microbiology*, *73*(16), 5261–5267. https://doi.org/10.1128/AEM.00062-07
